# The effects of denifanstat on hepatitis E virus replication and triggered inflammatory response

**DOI:** 10.1101/2025.10.16.682984

**Authors:** Jiahua Zhou, Maikel P. Peppelenbosch, Qiuwei Pan, Ibrahim Ayada

## Abstract

Metabolic dysfunction–associated steatotic liver disease (MASLD) is initiated by intra-cellular fatty acid accumulation, with macrophage-mediated inflammatory responses playing a pivotal role in driving disease progression. MASLD frequently coexists with viral infections, for example in the context of hepatitis E virus (HEV) infection which explores host lipid metabolism for its replication and secretion. But no targeted therapies currently exist for this complex comorbidity. Denifanstat, a fatty acid synthase (FASN) inhibitor developed for treating metabolic dysfunction-associated steatohepatitis (MASH), has shown to have antiviral activity against SARS-CoV-2 infection. In this study, we aim to assess the effects of denifanstat on HEV infection in the context of steatotic liver disease. We first employed human liver-derived primary organoids exposed to fatty acids and inoculated with HEV. Treatment with denifanstat had no major effect on HEV replication in the steatotic organoids. However, we found that denifanstat moderately inhibited viral replication in macrophages, and regulated HEV-triggered inflammatory responses. These results highlight the importance of understanding how the emerging MASH treatment may affect viral infections that coexist in patients.

## 1 Introduction

Metabolic dysfunction–associated steatotic liver disease (MASLD) is a leading cause of chronic liver disease worldwide. It is characterized by excessive lipid accumulation in the liver, leading to lipotoxicity and inflammation that can progress to metabolic dysfunction associated steatohepatitis (MASH), fibrosis, and hepatocellular carcinoma (HCC) [1]. In this pathogenic cascade, Kupffer cells (KCs), the liver-resident macrophages, play a pivotal role by releasing pro-inflammatory cytokines and chemokines in response to hepatic injury, thereby exacerbating inflammation and fibrosis [2].

The co-existence of MASLD and viral infections is clinically significant. For instance, individuals with HIV exhibit a higher prevalence of MASLD than the general population [3], while MASLD has been linked to worse outcomes in COVID-19 [4]. Chronic hepatitis C virus (HCV) infection is well-known to induce hepatic steatosis [5], and concurrent hepatitis B virus (HBV) infection may accelerate liver injury progression [6].

Denifanstat (TVB2640), an investigational small-molecule inhibitor of fatty acid synthase (FASN), represents a promising new treatment for MASH. FASN is a key enzyme in de novo lipogenesis, and its inhibition has shown efficacy in MASH, as evidenced by recent Phase 2b trials [7]. Intriguingly, denifanstat has been shown to potently inhibit SARS-CoV-2 infection in experimental models, attributed to its interference with lipid metabolism essential for viral replication [8]. Additionally, FASN modulates Toll-like receptor (TLR)-mediated macrophage activation and chemotaxis, suggesting that denifanstat could mitigate macrophage-driven inflammation in metabolic and infectious liver diseases [9].

In a Chinese cohort of acute hepatitis E virus (HEV), 50% had a steatotic liver while 24% was overweight and 9% obese [10]. Another European study found a higher anti-HEV IgG prevalence of 26% in biopsy-proven MASLD patients compared to 14% in healthy controls, however this was not independently associated with MASLD [11]. Interestingly, in this cohort presence of diabetes was independently associated with cirrhosis, which is inconsistent with the consensus. Therefore we here investigated the effects of denifanstat on HEV infection in a steatotic organoid model, and on viral replication and the triggered inflammatory responses in macrophages.

## 2 Methods

### 2.1 Reagents and preparation

Oleic acid (OA) and palmitic acid (PA) were obtained from Bio-Connect (The Netherlands). Denifanstat (TVB-2640; FASN-IN-2; ASC-40) was purchased from Sigma-Aldrich (The Netherlands). All compounds were dissolved in the optimal stock concentrations.

### 2.2 Cell culture and differentiation

Human adult intrahepatic cholangiocyte organoids (ICOs) were isolated and cultured following established protocols [12]. To maintain genetic stability and experimental reproducibility, only early-passage organoids (≤15 passages) were used in this study. THP-1 human monocytic cells were cultured in RPMI 1640 medium supplemented with 30 ng/mL of phorbol 12-myristate 13-acetate (PMA, Sigma-Aldrich Chemie BV) at 37°C for 48 hours to generate THP-1 macrophages.

### 2.3 HEV viruses and models

Plasmid constructs containing the full-length HEV genome (Kernow-C1 p6 clone; GenBank Accession Number JQ679013) and subgenomic GT3 HEV replicon coupled with a Gaussia luciferase reporter gene (p6Luc) were transcribed into genomic RNA in vitro from corresponding enzyme-digested and linearized plasmid DNA using mMessage mMachine T7 RNA kits (Invitrogen).

To establish HEV replication models, subgenomic GT3 HEV replicon RNA was electroporated into ICOs. For infectious virus production and model, full-length Kernow-C1 p6 genomic RNA was electroporated into Huh7 cells and ICOs.

For HEV-related Gaussia luciferase analysis (p6 Luc), the activity of secreted luciferase in the culture supernatant was measured by BioLux Gaussia Luciferase Flex Assay Kit (New England Biolabs, Ipswich, MA, USA).

### 2.4 Fatty acids exposure, viral inoculation, and denifanstat treatment

A 10x fatty acids (FA) stock solution was prepared by dissolving 250 uM OA and 125 uM PA in 5% bovine serum albumin (BSA). Three days post-electroporation, the organoid culture supernatant was removed and replaced with fresh medium containing FA and denifanstat.

THP-1 macrophages were washed twice with phosphate-buffered saline (PBS) to remove the residual reagents following differentiation. Subsequently, macrophages were inoculated with HEV particles (4.65 x 10^7 copy numbers/mL) and treated with denifanstat.

### 2.5 Quantification of viral replication and inflammatory gene expression

Viral RNA and inflammatory genes expression were quantified using SYBR Green-based quantitative real-time PCR (qRT-PCR) with Applied Biosystems™ SYBR Green PCR Master Mix (Thermo Fisher Scientific). GAPDH served as a housekeeping gene for normalizing target gene expression, employing the 2-(ΔΔCt) method.

### 2.6 Cell viability assays

For ICOs, the culture supernatant of organoids was discarded and then incubated with Alamar Blue (Life Technologies) at 37 °C with 5% CO2 for 2 hours. Next, Alamar Blue was collected for analyzing the metabolic activity of the organoids. Absorbance was determined by using fluorescence plate reader (CytoFluor Series 4000, Perseptive Biosystems) at the excitation of 530/25 nm and emission of 590/35 nm.

For macrophages, cell viability was determined by MTT assay (Sigma-Aldrich Chemie BV). Cells seeded in 48-well plates were incubated with 0.5 mg/mL MTT solution at 37°C with 5% CO2 for 4 hours. The medium was then removed, and 100 μL of DMSO was added to each well. The absorbance of each well was measured using a microplate absorbance reader (BIO-RAD) at wavelength of 570 nm.

### 2.7 Cytokine quantification by ELISA

The concentrations of IL-1β and TNF-a were measured with the Human IL-1 beta/IL-1F2 Quantikine ELISA Kit (Bio-Techne, Netherlands) and the Human TNF alpha Uncoated ELISA Kit (Thermo Fisher Scientific Life Sciences). Cytokine concentrations were calculated against standard curves generated for each assay.

### 2.8 Statistics

Statistical analysis was conducted using a non-paired, non-parametric test (Mann– Whitney U test; GraphPad Prism 8.0.2). All results were presented as mean ± standard deviation (SD), with statistical significance defined as *p < 0.05, **p < 0.01, and ***p < 0.001.

## 3 Results

### 3.1 Minimal effects of denifanstat treatment on HEV replication in organoid models

Subgenomic GT3 HEV replicon genomic RNA and full-length genomic RNA were electroporated into ICOs to establish replication and infectious models (Fig. 1A). FA exposure to mimic steatosis significantly inhibited HEV-replication related luciferase activity in the subgenomic model (p < 0.001; Fig. 1B). However, treatment with denifanstat had no significant effects on viral replication in organoids with or without FA exposure (Fig. 1C-1H).

**Figure 1.**
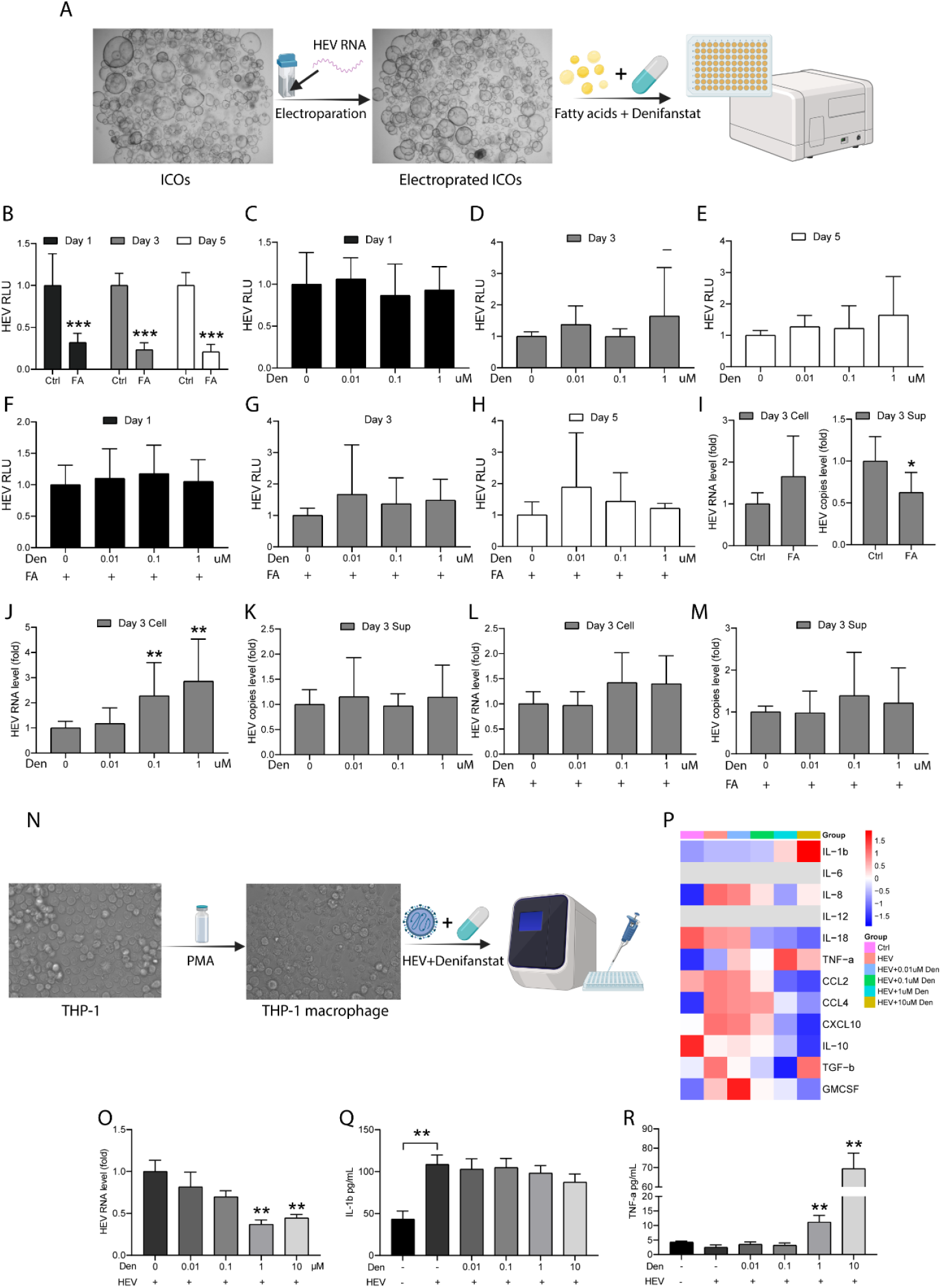
Denifanstat treatment on HEV infection in human liver organoids and macrophages. A, Schematic overview of electroporating ICOs with HEV genomic RNA, and subsequent exposure of fatty acids (FA) and treatment with denifanstat. The schematic illustration was generated in Biorender. B-H, Exposure with or without fatty acids and treatment with denifanstat in ICOs with p6-Luc HEV replicon model. Luciferase activity in ICOs from day 1, 3 and 5 post-electroporation (n = 6) I-M, Exposure with or without fatty acids and treatment with denifanstat in ICOs based p6 HEV infectious model. HEV RNA level in organoid lysate and virus copies in supernatant from ICOs on day 3 post-electroporation (n = 6) N, Schematic overview of polarizing THP-1 macrophages and subsequent inoculation with HEV and treatment of denifanstat. The schematic illustration was generated in Biorender. O, HEV RNA levels in lysate of macrophages after HEV inoculation and treatment by denifanstat for 2 days (n = 6). P, The mRNA expression of inflammatory genes in macrophages after HEV inoculation and treatment by denifanstat for 2 days (n = 6). Q,R IL-1β and TNF-α protein level were quantified by ELISA in macrophages after HEV inoculation and treatment by denifanstat for 2 days (n = 6). *p < 0.05, **p < 0.01, and ***p < 0.001.

In organoids harboring infectious HEV, FA exposure slightly reduced the amount of released viruses (p < 0.05; Fig. 1I). In organoids without FA exposure, treatment with denifanstat at 0.1 µM and 1 µM moderately elevated the cellular levels of HEV RNA (Fig. 1J), but not on the level of released viruses (Fig. 1K). No significant effect on HEV RNA level was observed in organoids with FA exposure (Fig. 1L and 1M). Of note, FA exposure had some inhibitory effects on organoid growth (Supplementary Fig. 1A).

### 3.2 Denifanstat moderately inhibits HEV replication and regulates inflammatory responses in macrophages

Human THP-1 monocytic cells were differentiated into macrophages through 48-hour PMA exposure. Next, we inoculated macrophages with HEV and treated with denifanstat for 48 h (Fig. 1N). Our results demonstrated that denifanstat exhibited minor inhibitory effects on macrophage growth (Supplementary Fig. 1B), but significantly inhibited HEV replication in a dose-dependent manner, with 1 µM concentration achieving over 50% inhibition (p < 0.01; Fig. 1 O).

HEV triggered robust inflammatory responses in macrophages, leading to upregulation of several inflammatory genes (Fig. 1P). Remarkably, denifanstat treatment markedly suppressed the expression of most inflammatory genes at mRNA levels, except for IL-1β and TNF-α (Fig. 1P). At the protein levels, denifanstat had no major effect on IL-1β (Fig. 1Q), but significantly elevated TNF-α production by approximately 14-fold at 10 µM concentration (p < 0.01; Fig. 1R).

## 4 Discussion

Denifanstat specifically targets and inhibits FASN, the key enzyme responsible for fatty acid synthesis. It exhibits antiviral potential primarily by disrupting host lipid metabolism, thereby interfering with viral replication and release, such as in SARS-CoV-2. The denifanstat concentrations that we used align with a previous study tested in SARS-CoV-2 experimental models [8], and are also preclinically achievable concentrations [13]. However, we did not observe notable antiviral activity of denifanstat again HEV infection in human liver organoids with or without FA exposure. Nevertheless, targeting lipid metabolism has been shown to inhibit HEV maturation or membrane association by depleting intracellular lipids, trapping viral particles within cells and preventing efficient egress [14].

Previous studies have shown that HEV can infect macrophages and trigger inflammatory responses [15]. Additionally, hepatic macrophages contribute to MASH progression by secreting pro-inflammatory cytokines [2]. In our study, we observed that denifanstat moderately inhibits HEV replication in macrophages and broadly regulated the expression of HEV-triggered inflammatory genes. In inflammatory macrophages, the tricarboxylic acid (TCA) cycle is known to be disrupted by following citrate and succinate accumulation which are related to biosynthesis of fatty acids. Inhibition of FASN disrupts membrane composition, impairs cholesterol retention, and alters Rho GTPase trafficking, ultimately dampening inflammatory responses [16].

In summary, denifanstat had no major effect on HEV replication in organoids, but moderately inhibited the replication in macrophages, as well as modulated inflammatory responses. Thus, the mode-of-action of denifanstat could be cell type specific. Given the relatively common co-existence of steatotic liver disease and viral infections, the currently witnessed significant activity of the drug development pipeline for MASH, should also consider studying the effects of these drugs on both conditions.

## Supplimentary Fig

**Fig. S1.**
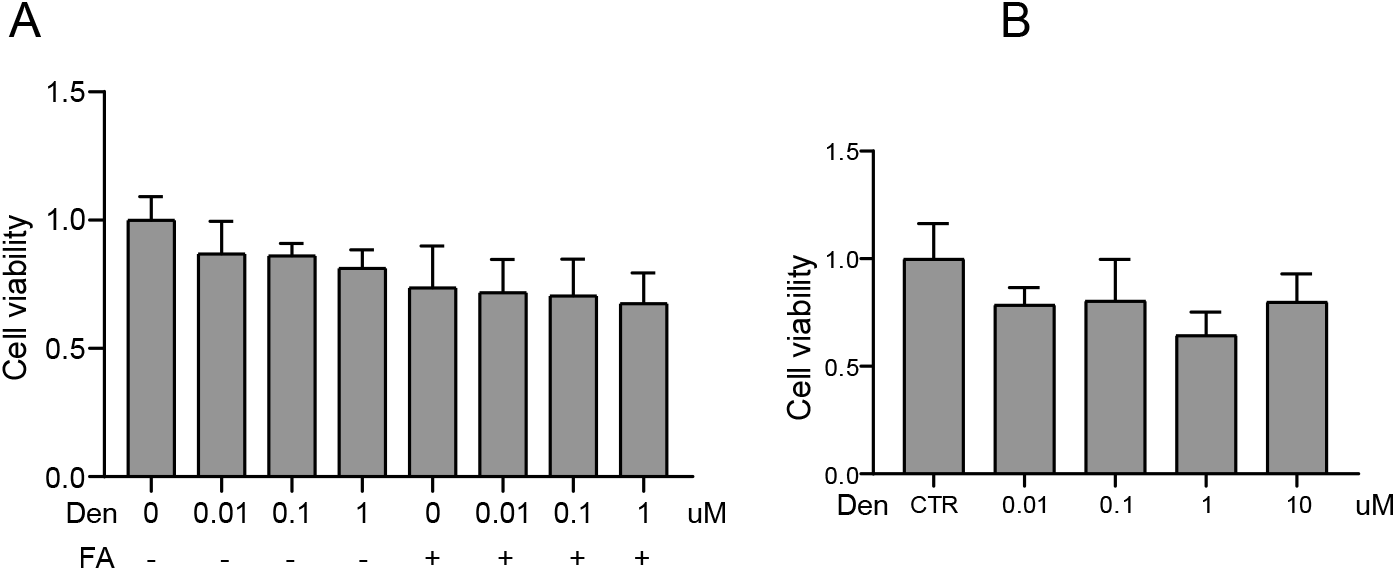
Treatment on cell viability. A, Exposure with fatty acids (FA) and treatment with denifanstat (Den) in ICOs with p6-Luc HEV replicon model. Cell viability of ICOs from day 3 post-electroporation (n = 6) B, Cell viability of THP-1 macrophages after HEV inoculation and treatment by denifanstat for 2 days (n = 6)

